# Intratumoral dose heterogeneity promotes adaptive anti-tumor immunity and predicts clinical response to radiopharmaceutical therapy

**DOI:** 10.64898/2026.07.23.740178

**Authors:** Maya E. Takashima, Ohyun Kwon, Zachary Ells, Vincent R. Li, Charlotte Sawicki, Rene Welch Schwartz, Shin Hye Ahn, Meredith Hyun, Malick Bio Idrissou, Tracy J. Berg, Paul A. Clark, Michael Lawless, Abby Besemer, Tyler Bradshaw, Scott Perlman, Wonjong Jin, Michael Antonelli, Adedamola O Adeniyi, Campbell Donnelly Haasch, Tessa Chen, Yujuan Wang, Ria Kumari, Reinier T. Hernandez, Jamey Weichert, Anthony P. Belanger, Irene Ong, John Floberg, Catherine Meyer, Amar U Kishan, Jeremie Calais, Bryan Bednarz, Zachary S. Morris

**Affiliations:** Department of Human Oncology, University of Wisconsin School of Medicine and Public Health, University of Wisconsin-Madison, Madison, WI, USA 53705; Department of Medical Physics, University of Wisconsin School of Medicine and Public Health, University of Wisconsin-Madison, Madison, WI, USA 53705; Department of Nuclear Medicine and Theranostics, Ahmanson Translational Theranostics Division, David Geffen School of Medicine, UCLA, Los Angeles, California, USA; Department of Biostatistics and Medical Informatics, University of Wisconsin School of Medicine and Public Health, University of Wisconsin-Madison, Madison, WI, USA 53705; Molecular Cancer Imaging Facility, Department of Imaging, Dana-Farber Cancer Institute, Boston, Massachusetts, USA; Department of Radiology, Harvard Medical School, Boston, Massachusetts, USA; Department of Radiology, University of Wisconsin School of Medicine and Public Health, University of Wisconsin-Madison, Madison, WI, USA 53705; Department of Radiation Oncology, UCLA, Los Angeles, California, USA

## Abstract

Radiopharmaceutical therapies (RPT) deliver non-uniform radiation dose in tumors and the impact of this on response is poorly understood. Dose heterogeneity could engender treatment resistance in low dose regions, yet we hypothesize that a broader array of dose-dependent immuno-radiobiological mechanisms in tumor microenvironments (TME) and preservation of immune function in low-dose regions could promote adaptive anti-tumor immunity and response. In murine models, non-uniform lutetium-177 delivering <2.5 Gy to >20 Gy in a TME induced broader immunomodulatory effects and T cell-dependent survival improvement compared to more uniform distributions. Preserving low-dose regions promoted dendritic cell activation and TME infiltration of clonally expanded CD8^+^ T cells. In three independent cohorts of patients with prostate cancer, heterogeneous tumor dose distribution strongly correlated with improved clinical outcomes. These findings defy expected radiobiological dose-response and define a novel mechanism of action for RPT, supporting clinical investigation of dose distribution for optimizing patient selection and personalized dosing.

## INTRODUCTION

Radiation therapy (RT) activates distinct, dose-dependent immune responses within the tumor microenvironment (TME) (1). Low-dose RT promotes immune trafficking, moderate-dose RT optimally induces a type I interferon response, and high-dose RT triggers immunogenic cell death and upregulation of susceptibility markers, including Major Histocompatibility Complex Class I (MHCI) and FAS (2–7). Traditional RT aims to maximize uniform dose coverage across a target volume to avoid resistance due to under treatment (8). However, emerging evidence suggests that deliberate dose heterogeneity, combining low, moderate, and high dose regions within a single TME, maximizes anti-tumor immunity (9–17). Although these methods could achieve a heterogeneous dose in a single tumor, in settings of widespread metastatic cancers there is a limitation to the number of tumor sites that can feasibly and safely be treated by brachytherapy or spatially fractionated RT (SFRT).

Radiopharmaceutical therapy (RPT) can deliver radiation selectively to disseminated metastatic tumor sites (18–20). RPT characteristically delivers a highly heterogeneous dose distribution within the TME due to non-uniform tumor perfusion and variation in distribution and expression of the RPT-targeted molecule in the TME. In the broader context of molecularly targeted therapeutics, such heterogeneity is a primary driver of resistance as cells lacking a drug target evade treatment entirely. Under conventional radiobiology paradigms, RPT dose heterogeneity is viewed as a potential pitfall that may limit therapeutic efficacy, particularly for agents delivering isotopes like ^177^Lu, with a mean range in tissue that is short (0.67 mm) relative to the size of a typical TME. Yet clinical activity has been established for such agents, suggesting that it may be valuable to further assess the mechanistic underpinnings of response to RPT (21, 22). Furthermore, RPT is typically prescribed at a fixed injected activity, which is simple but leads to large variation in the mean absorbed tumor dose between patients. In the absence of dosimetry, the actual dose of RT delivered to tumors in a specific patient remains unknown; however, many clinical centers are increasingly applying imaging-based dosimetry to predict absorbed dose to the tumor and normal tissues from RPT (23, 24). Few preclinical or clinical studies have investigated the dose- and dose heterogeneity-dependent immuno-radiobiological effect of RPT in the TME. Clarifying these mechanistic relationships is critical to capitalize on tumor dosimetry for personalized approaches to prescribing RPT.

In this manuscript, heterogeneous refers to a dose distribution that contains a range from therapeutically low (< 2.5 Gy) to high (>20 Gy) dose in a TME while more homogeneous distributions exhibit a narrower range. To evaluate the effects of RPT dose heterogeneity on the anti-tumor immune response, we utilize two ^177^Lu-labeled agents: ^177^Lu-PNT6555, targeting fibroblast activation protein alpha (FAP-α) in engineered homogeneous versus heterogeneous tumor models (25), and ^177^Lu-NM600, an alkylphosphocholine analog that selectively targets tumor cells over endogenous stroma (26). We hypothesize that for a given mean tumor absorbed dose from lutetium-177, a heterogeneous as compared to a more homogeneous dose distribution will more effectively activate multiple dose-dependent immuno-radiobiologic mechanisms and enhance tumor response through adaptive anti-tumor immunity. Our preclinical findings corroborate this hypothesis, and the translational relevance of these findings is supported by an exploratory and confirmatory assessment of intratumoral heterogeneity and response to PSMA-targeting lutetium-177 among multiple independent cohorts of men with metastatic prostate cancer.

## RESULTS

### Microscale dosimetry quantifies dose heterogeneity from lutetium-177

To evaluate RPT dose heterogeneity, we utilized two syngeneic murine models: a B16 melanoma model with controlled target heterogeneity (engrafted as either 100% mFAP^+^ cells or a mixed mFAP^+^/mFAP^-^ population) targeted by ^177^Lu-PNT6555 and a KP sarcoma model with endogenous stromal heterogeneity targeted by ^177^Lu-NM600. Serial SPECT/CT imaging combined with Monte Carlo dosimetry established macroscopic mean tumor absorbed doses (Figure 1A and 1B)(27). High-resolution *ex vivo* imaging with the ionizing radiation Quantum Imaging Detector (iQID) subsequently mapped microscale dose distributions and dose-volume histograms (DVHs) (Figure 1C-1F)(28). At an equivalent mean tumor dose of 5 Gy, homogeneous B16-mFAP^+^ tumors yielded a narrow dose distribution (V_2.5 Gy_= 90% with a max dose of 10.83 Gy; Figure 1C and 1E). In contrast mixed B16-mFAP^+^/mFAP^-^ tumors exhibited pronounced dose heterogeneity (V_2.5 Gy_=75% with a max dose of 23.43 Gy; Figure 1C and 1E). Similarly, KP Sarcoma tumors delivered a broad spectrum of low, moderate, and high doses (V_2.5 Gy_=74% with a max dose of 25.97 Gy; Figure 1D and 1F). These findings demonstrate that partial target expression or stromal architecture creates substantial microscale dose heterogeneity, delivering high-dose peaks alongside low-dose regions within a single tumor even at low mean tumor doses.

**Figure 1.**
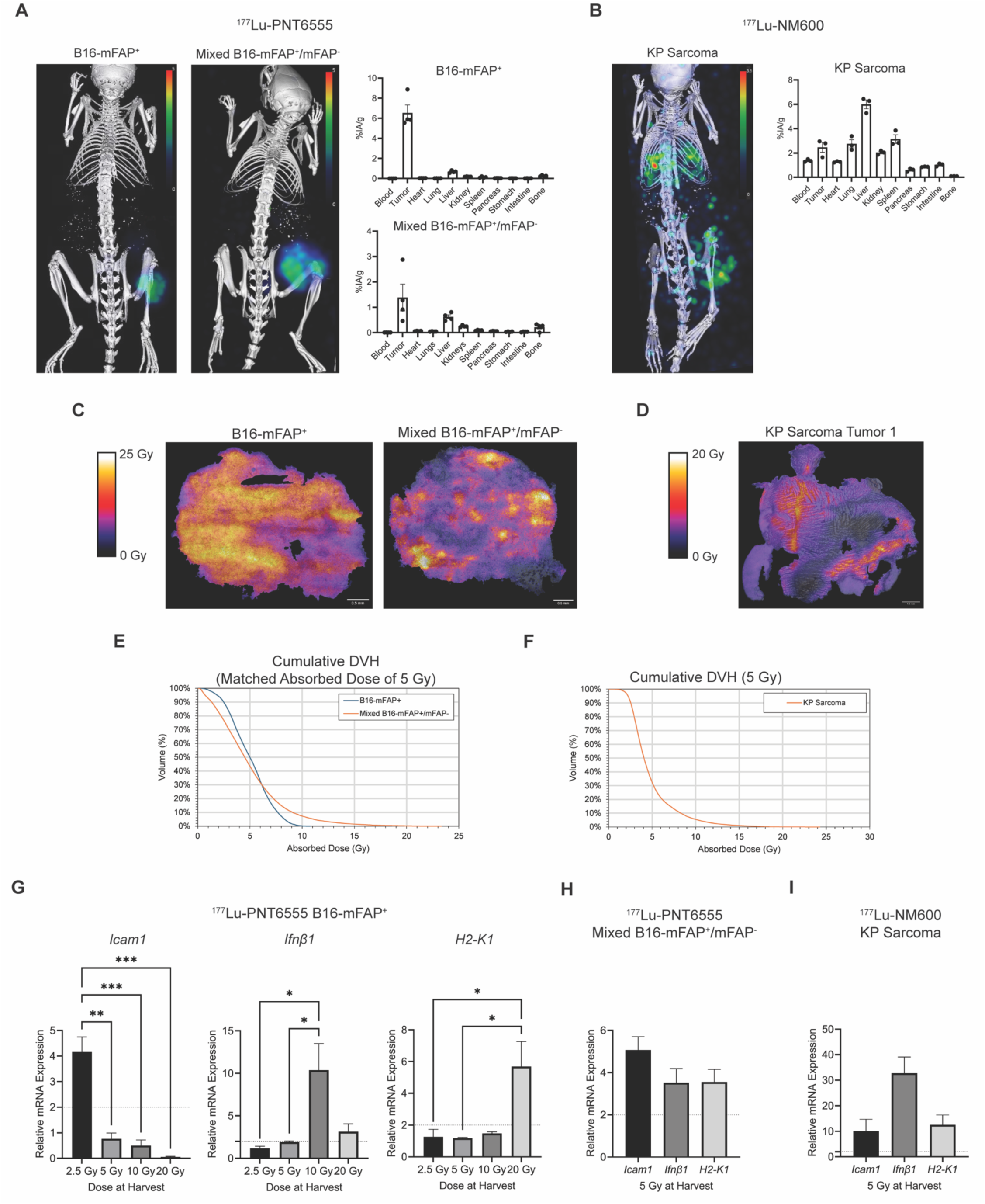
Microscale dosimetry quantifies dose heterogeneity from Lutetium-177, driving concurrent induction of dose-dependent immune-related gene expression. A-F) *in vivo* and *ex vivo* imaging for ^177^Lu-PNT6555 and ^177^Lu- NM600 A-B) Maximum intensity projections (MIPs) of SPECT/CT images at 24h post injection of 500 μCi of A) B16-mFAP+ (left) and mixed B16-mFAP+/mFAP- (right) melanoma tumor-bearing mice with ^177^Lu-PNT6555, B) KP Sarcoma tumor-bearing mice with ^177^Lu-NM600. C) Absorbed dose maps with isodose lines from ionizing quantum imaging detector images for B16-mFAP^+^ (left) and mixed B16-mFAP^+^/mFAP^-^ (right) at 24h post injection of 666 μCi ^177^Lu-PNT6555. D) Absorbed dose map with isodose lines from iQID for KP Sarcoma at 24h post injection of 250 μCi ^177^Lu-NM600. E) Cumulative DVH curves for a B16-mFAP^+^ (blue) and mixed B16-mFAP^+^/mFAP^-^ (orange) tumor receiving a matched absorbed dose of 5 Gy form ^177^Lu-PNT6555. F) Cumulative DVH curves for two KP Sarcoma tumors receiving matched absorbed dose of 5 Gy from ^177^Lu-NM600. G-I) B16-mFAP^+^, mixed B16-mFAP^+^/mFAP^-^, or KP Sarcoma tumor-bearing mice were treated with ^177^Lu-PNT6555 or ^177^Lu-NM600 on day 0 and harvested at day when tumors received 2.5, 5. 10, or 20 Gy. qPCR was used to quantify gene expression of *Icam1*, *Ifnβ1*, *H2-K1* and reported as fold change normalized to untreated control. G) B16-mFAP^+^ tumor-bearing mice treated with ^177^Lu- PNT6555. H) Mixed B16-mFAP^+^/mFAP^-^ tumor-bearing mice treated with ^177^Lu-PNT6555. I) KP Sarcoma tumor-bearing mice treated with ^177^Lu-NM600. G-I) N=4 all groups. One-way ANOVA was used to compare fold change in expression between groups.

### Heterogeneous RPT dose activates expression of immune susceptibility markers with diverse dose-response profiles in a single TME

To investigate effects of RPT on immune-related mechanisms that exhibit distinct dose-response profiles, we measured *Icam*, *Ifnβ1*, and *H2-K1* in mice bearing B16-mFAP^+^ tumors treated across a range of ^177^Lu-PNT6555 mean doses (2.5, 5, 10, or 20 Gy by day 7). RT-qPCR demonstrated distinct dose-dependent peaks (p<0.05): *Icam* peaked at 2.5 Gy (low dose), *Ifnβ1* peaked at 10 Gy (moderate dose), and *H2-K1* at 20 Gy (high dose; Figure 1G), consistent with dose-dependent effects observed in EBRT (9). We then tested whether non-uniform RPT could concurrently activate all three immune mechanisms within a single TME. In mixed B16- mFAP^+^/mFAP^-^ tumors treated with a 5 Gy mean dose from ^177^Lu-PNT6555, microscale dose heterogeneity induced robust simultaneous expression of *Icam, Ifnβ1*, and *H2-K1* relative to controls (Figure 1H). This concurrent activation was confirmed in KP sarcoma tumors receiving 5 Gy from ^177^Lu-NM600 (Figure 1I). Thus, non-uniform lutetium-177 can activate multiple dose-dependent immune-related mechanisms in a single TME.

### High intrinsic heterogeneity from RPT that achieves low-, moderate-, and high-dose regions leads to better overall survival

We evaluated whether non-uniform RPT enhances therapeutic response compared to a uniform distribution using B16-mFAP^+^ (low target heterogeneity) and mixed B16-mFAP^+^/mFAP^-^ (high target heterogeneity) melanoma models (Table 1 and Figure S1). To achieve an equivalent mean tumor dose of 5 Gy, mixed tumors required higher injected activity, establishing a broad dose distribution spanning low, moderate, and high dose regions whereas uniform tumors receive a narrow distribution ranging only from low to moderate doses. At this equivalent 5 Gy mean tumor dose, mixed B16-mFAP^+^/mFAP^-^ tumors exhibit significantly improved tumor control and overall survival compared to B16-mFAP^+^ tumors (p = 0.0125; Figure 2A and 2B). These data indicate that for a given mean tumor dose, therapeutic efficacy is maximized when dose variance is elevated to include low, moderate, and high dose regions within the TME.

**Figure 2.**
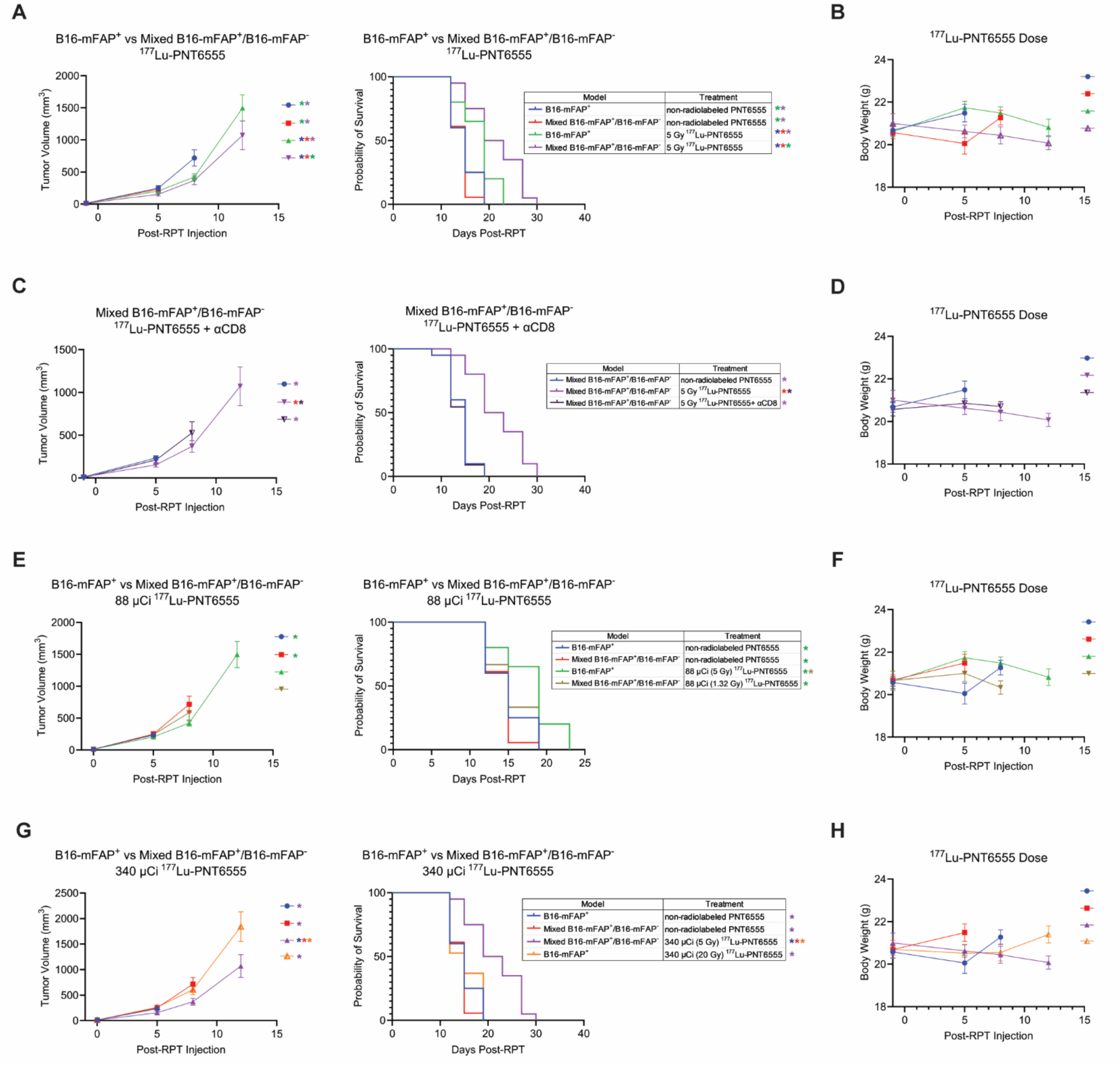
Increased radiation dose heterogeneity from ^177^Lu-PNT6555 improve overall survival. A-H) B16- mFAP^+^ or mixed B16-mFAP^+^/mFAP^-^ tumor-bearing mice were randomized to non-radiolabeled PNT6555, 1.32, 5, 20 Gy ^177^Lu-PNT6555 on day 0. A-B) Effects of more heterogeneous compared to less heterogeneous 5 Gy ^177^Lu- PNT6555. C-D) CD8 depletion with ^177^Lu-PNT6555 in mixed B16-mFAP^+^/mFAP^-^. E-H) Effects of matched activity from ^177^Lu-PNT6555. A-H) Injected activities were as followed: B16-mFAP^+^ 5 Gy ^177^Lu-PNT6555=88 µCi; B16- mFAP^+^ 20 Gy ^177^Lu-PNT6555=340 µCi; mixed B16-mFAP^+^/mFAP^-^ 5 Gy ^177^Lu-PNT6555=340 µCi; mixed B16- mFAP^+^/mFAP^-^ 1.32 Gy=88 µCi. Linear mixed model was used to compare tumor growth and log-rank test was used to compare survival. *p < 0.05 (A, C, E, G), by Kaplan-Meier method (A, C, E, G) the color of the asterisk represents the group from which the group differs.

**Table 1.** Injected RPT activity for tumor dose prescriptions.

| Tumor Model | RPT Agent | Tumor Absorbed Dose Prescription (Gy) | Injected Activity ($\mu\text{Ci}$) |
| --- | --- | --- | --- |
| B16-mFAP | $^{177}\text{Lu}$-PNT6555 | 5 | 88 |
| | $^{177}\text{Lu}$-PNT6555 | 10 | 175 |
| | $^{177}\text{Lu}$-PNT6555 | 20 | 340 |
| | $^{177}\text{Lu}$-PNT6555 | 40 | 702 |
| B16-mixed | $^{177}\text{Lu}$-PNT6555 | 1.32 | 88 |
| | $^{177}\text{Lu}$-PNT6555 | 5 | 340 |
| KP sarcoma | $^{177}\text{Lu}$-NM600 | 2.5 | 125 |
| | $^{177}\text{Lu}$-NM600 | 5 | 251 |
| | $^{177}\text{Lu}$-NM600 | 10 | 502 |
| | $^{177}\text{Lu}$-NM600 | 15 | 754 |

Given that heterogeneous external beam and brachytherapy can enhance anti-tumor immunity (9, 11, 12, 29), and low mean tumor doses promote immune activation (30, 31), we hypothesized that CD8 T cells mediate the efficacy of heterogeneous RPT. Depletion of CD8 T cells during 5 Gy ^177^Lu-PNT6555 treatment in mixed B16-mFAP^+^/mFAP^-^ tumors significantly accelerated tumor growth compared to non-depleted controls p<0.0001; Figure 2C and 2D), whereas depletion had no effect on non-radiolabeled controls (Figure S2A and S2B). These results demonstrate that CD8 T cells are required for the enhanced therapeutic efficacy driven by heterogeneous RPT.

To isolate the effect of dose distribution from total injected activity, we performed activity matched control experiments (Table 1). Administering 88 µCi to mixed B16- mFAP^+^/mFAP^-^ tumors (the activity delivering 5 Gy in B16-mFAP+ tumors) failed to suppress tumor growth or improve survival (Figure 2E and 2F). Conversely, administering 340 µCi to B16-mFAP^+^ tumors (the activity delivering 5 Gy in mixed tumors) yielded no therapeutic benefit (Figure 2G and 2H). This demonstrates the complexity of adequately modeling RPT dose-response relationships using conventional radiobiology and optimally dosing RPT. Given the distinct dose-response relationships of lutetium-177, we next evaluated the use of heterogeneity metrics to predict response.

### Tumor dose heterogeneity metrics predict response to lutetium-177

Given the role of CD8 T cells in mediating RPT efficacy, we evaluated how off-target radiation impacts systemic immune cell counts, as lymphopenia is a known negative predictor of radiation response (32–35). We hypothesized that optimal RPT dosing requires the highest dose that preserved dose heterogeneity (did not eliminate low-dose intratumoral regions) while avoiding systemic lymphopenia. In B16-mFAP^+^ tumor-bearing mice treated with 0, 5, 10, or 20 Gy from ^177^Lu-PNT6555 (Table 1), peripheral lymphocytes counts at day 14 were significantly reduced at 20 Gy compared to controls (p= 0.0068; Figure 3A). Cumulative DVH curves demonstrated that 10 Gy uniquely achieved high-dose in regions of a TME while also preserving low-dose regions in this TME (Figure 3B). Consequently; 10 Gy from ^177^Lu-PNT6555 significantly prolonged overall survival compared to 5 Gy (p<0.0001; Figure 3C and 3D).

**Figure 3.**
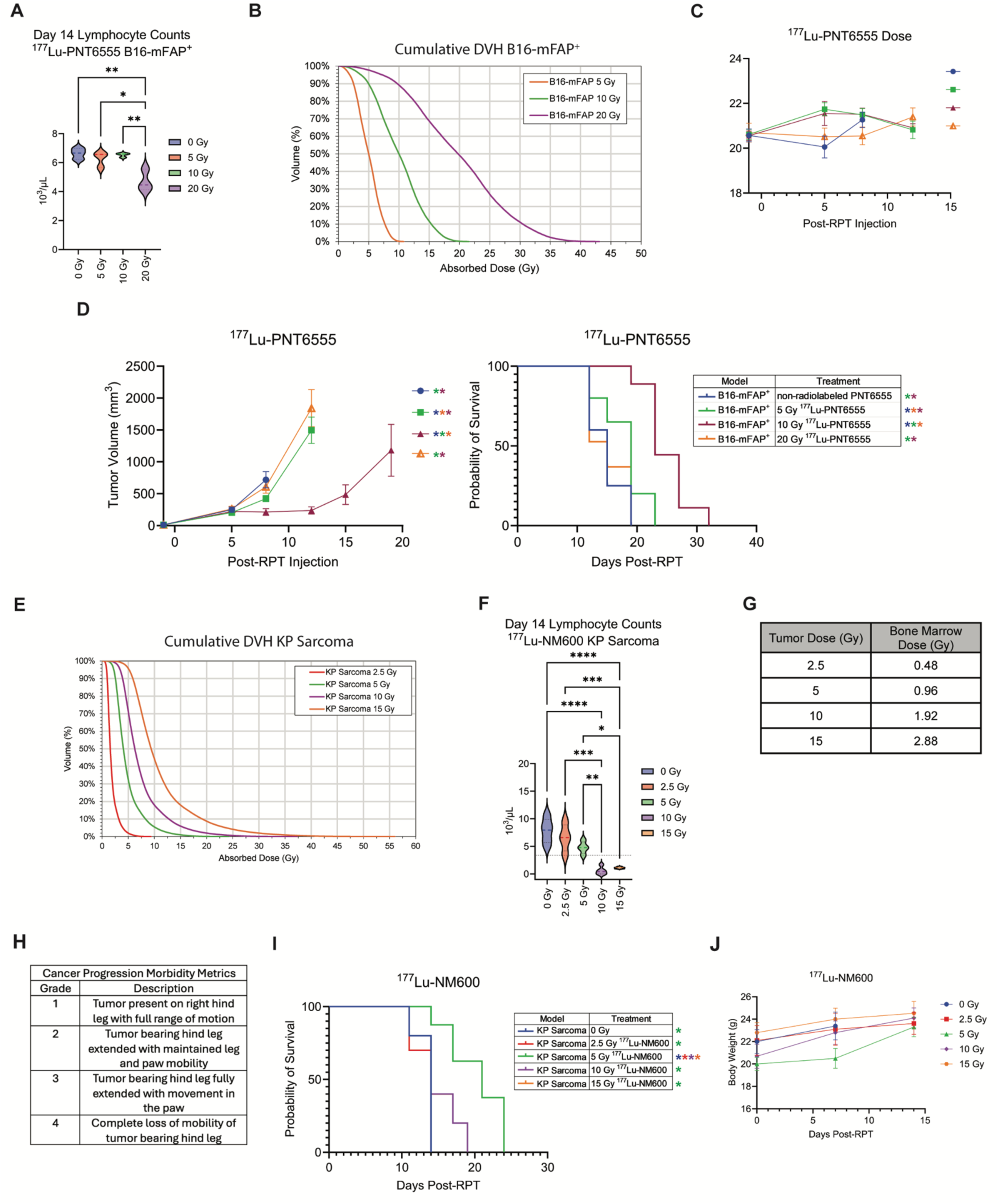
Radiation dose heterogeneity from ^177^Lu-PNT6555 and ^177^Lu-NM600 including a low, moderate, and high dose region improves overall survival. A-D) B16-mFAP^+^ tumor-bearing mice were randomized to non-radiolabeled PNT6555, 5, 10 Gy, 20 Gy ^177^Lu-PNT6555 on day 0. A) Lymphocyte counts at day 14 following injection of ^177^Lu-NM600. B) Cumulative DVH curves for B16-mFAP^+^ tumors receiving 5, 10, 20 Gy ^177^Lu-PNT6555. C) Overall mouse body weights during survival study. D) Effects of different mean tumor dose from ^177^Lu-PNT6555. A-D) Injected activities were as followed: B16-mFAP^+^ 5 Gy ^177^Lu-PNT6555=88 µCi; B16-mFAP^+^ 10 Gy ^177^Lu- PNT6555=175 µCi; B16-mFAP^+^ 20 Gy ^177^Lu-PNT6555=340 µCi. E-J) KP sarcoma tumor-bearing mice were randomized to untreated (0 Gy), 2.5, 5. 10, 15 Gy ^177^Lu-NM600 on day 0. E) Cumulative DVH curves for KP sarcoma tumors receiving 2.5, 5, 10, 15 Gy ^177^Lu-NM600. F) Lymphocyte counts at day 7 and 14 following injection of ^177^Lu- NM600. G) Mean bone marrow dose from ^177^Lu-NM600 corresponding to mean tumor dose. H) Cancer progression morbidity metrics used to determine endpoint for KP sarcoma bearing mice. I) Effects of ^177^Lu-NM600 doses on overall survival. J) Overall mouse body weights during survival study. E-J) Injected activities were as followed: KP Sarcoma 2.5 Gy ^177^Lu-NM600=125 µCi; KP Sarcoma 5 Gy ^177^Lu-NM600=251 µCi; KP sarcoma 10 Gy ^177^Lu- NM600=502 µCi; KP Sarcoma 15 Gy ^177^Lu-NM600=702 µCi. B-G) N=20: B16-mFAP^+^ non-radiolabeled PNT6555, mixed B16-mFAP^+^/mFAP^-^ non-radiolabeled PNT6555, B16-mFAP^+^ 5 Gy ^177^Lu-PNT6555, B16-mFAP^+^ 5 Gy ^177^Lu- PNT6555 + αCD8, B16-mFAP^+^ 20 Gy ^177^Lu-PNT6555, mixed B16-mFAP^+^/mFAP^-^ 5 Gy ^177^Lu-PNT6555, mixed B16- mFAP^+^/mFAP^-^ 1.32 Gy; n=10: mixed B16-mFAP^+^/mFAP^-^ 5 Gy ^177^Lu-PNT6555 + αCD8, B16-mFAP^+^ 10 Gy ^177^Lu- PNT6555. K) n=10: KP Sarcoma 0 Gy, 2.5 Gy ^177^Lu-NM600, 5 Gy ^177^Lu-NM600, 10 Gy ^177^Lu-NM600, and 15 Gy ^177^Lu-NM600. Linear mixed model was used to compare tumor growth and log-rank test was used to compare survival. *p < 0.05. **p < 0.01, ***p < 0.001, ****p < 0.0001 by one-way ANOVA (A and E). *p < 0.05 by linear mixed model (D), by Kaplan-Meier method (D and H) the color of the asterisk represents the group from which the group differs.

To test the generalizability of this framework, we evaluated SvJ mice bearing KP sarcoma treated with ^177^Lu-NM600 at 2.5, 5, 10, or 15 Gy (Table 1; Figure 3E). While all doses significantly decreased blood lymphocyte counts at day 7, counts recovered by day 14 in the 2.5 and 5 Gy groups but remained significantly depleted at 10 Gy (p<0.0001) and 15 Gy (p<0.0001; Figure 3F). To establish a systemic dosimetry parameter relevant to the anti-tumor immune response, we evaluated the relationship between mean tumor dose and mean bone marrow dose. Mean bone marrow dose increased proportionally with mean tumor dose (0.48 Gy bone marrow at 2.5 Gy tumor; 0.96 Gy at 5 Gy; 1.92 Gy at 10 Gy; 2.88 Gy at 15 Gy; Figure 3G). Systemic lymphopenia occurred when mean bone marrow dose exceeded 1.92 Gy, aligning with the accepted 2 Gy mean bone marrow limit (36). For KP sarcoma, 5 Gy from ^177^Lu-NM600 preserved low-dose tumor regions while keeping bone marrow dose below 1 Gy. Following a 4- grade leg morbidity scale, mice were euthanized upon reaching Grade 4 impairment (Figure 3H). Treatment with 5 Gy from ^177^Lu-NM600 significantly extended overall survival to 21 days compared to 0 Gy (14 days, p=0.004), 2.5 Gy (14 days, p=0.003), 10 Gy (14 days, p=0.046), and 15 Gy (14 days, p=0.046), without impacting body weight (Figure 3I and 3J). Together, these data are consistent with the hypothesis that optimal RPT efficacy is achieved at the highest injected activity that still preserves low-dose regions in this TME and that keeps mean bone marrow dose below 2 Gy to prevent systemic lymphopenia.

### Heterogeneous dose of RPT promotes adaptive anti-tumor immunity

To investigate whether non-uniform RPT promotes adaptive anti-tumor immunity, we evaluated dendritic cell (DC) activation and T cell dynamics in tumors and tumor-draining lymph nodes (TDLNs) following 5 Gy mean tumor dose from ^177^Lu-PNT6555 in uniform (B16-mFAP^+^) and heterogeneous (mixed B16-mFAP^+^/mFAP^-^) models (Figure 4 and S4).

**Figure 4.**
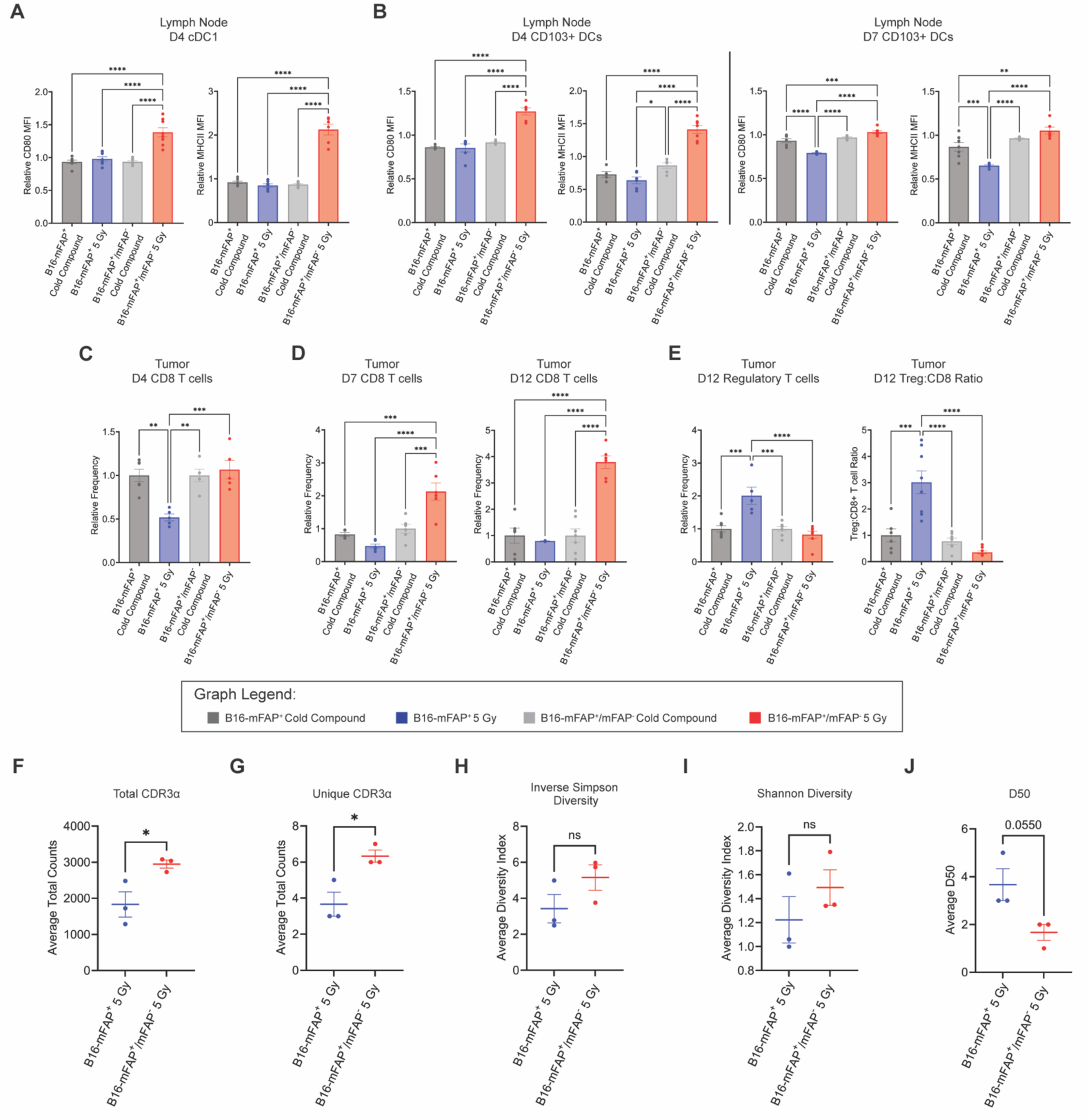
Radiopharmaceutical therapy dose heterogeneity including a low, moderate, and high dose region promotes adaptive immunity through dendritic cell activation in the tumor draining lymph node and T cell infiltration and clonal expansion in the tumor. A-E) B16-mFAP^+^ or mixed B16-mFAP^+^/mFAP^-^ tumor-bearing mice were treated with non-radiolabeled PNT6555 (Cold Compound) or 5 Gy ^177^Lu-PNT6555 on day 0 and tumors and tumor draining lymph nodes were harvested at day 4, 7, and 12. Results shown as relative frequencies and MFIs to non-radiolabeled PNT6555 (Cold Compound). A) Analysis of CD80 and MHC class II MFI in tumor draining lymph node (TDLN) type 1 conventional dendritic cells, cDC1 (CD45+CD11b+CD11c+MHCII+XCR1+) at day 4. B) Analysis of CD80 and MHC class II MFI in TDLN CD103+ DCs (CD45+CD11b+CD11c+CD103+) at day 4 and 7. C) Analysis of CD8^+^ T cells (CD45+CD3+CD8+) preservation in tumor at day 4. D) Analysis of CD8+ T cell infiltration in tumor at day 7 and 12. E) Analysis of Regulatory T cell (CD45+CD3+CD4+FoxP3+) infiltration and the Regulatory T cell to CD8+ T cell ratio in tumor at day 12. F-J) TCR sequencing of RNA extracted from B16-mFAP^+^ or mixed B16-mFAP^+^/mFAP^-^ tumors. F) Total number of CDR3α sequences. G) the counts of the unique CDR3α sequences. H) the inverse Simpson diversity index. I) the Shannon diversity index. J) the D50 value. n = 3 mice per group. A-E) *p < 0.05. **p < 0.01, ***p < 0.001, ****p < 0.0001 by one-way ANOVA. F-J) *p<0.05 by unpaired T-Test.

We first assessed the TDLN, where antigen presentation and naïve T cell priming occur (37). Heterogeneous RPT significantly elevated key activation markers (CD80, MHCII) on primary CD8-priming DC subsets (38), including cDC1s at day 4 (p<0.0001) and CD103+ migratory DCs at days 4 and 7 (p<0.0001; Figure 4A and 4B). Significant increases in activation markers were also observed across cDC2 and pDC populations in the TDLN (Figure S3). Preserving low-dose TME regions thus maintains functional DC populations required for T cell priming in TDLN.

In the tumor, where radiosensitive lymphocytes are vulnerable to radiation (39), non- uniform RPT preserved and recruited effector populations (9, 11, 12, 29). While uniform RPT depleted intratumoral CD8 T cells at day 4 (p=0.0007), heterogeneous RPT significantly increased CD8 T cell frequencies at days 7 (p<0.0001) and 12 (p<0.0001; Figure 4C and 4D). Furthermore, uniform RPT significantly increased the immunosuppressive regulatory T cell to CD8 T cell ratio compared to heterogeneous RPT (p<0.0001; Figure 4E). Corresponding increases in MHCI expression and CD4 T cell infiltration are detailed in Figure S3.

To evaluate the clonal dynamics of CD8 T cells in the TME following RPT, we performed bulk TCR deep sequencing on tumor-infiltrating lymphocytes 12 days post-treatment (non-radiolabeled conditions shown in Figure S5). Heterogeneous RPT significantly increased both the number of unique CDR3α sequences (p=0.0232) and total CDR3α sequence counts (p=0.0381) compared to uniform RPT (Figure 4F and 4G). While overall clonal diversity metrics (Shannon and inverse Simpson indices) were unchanged (Figure 4H and 4I), heterogeneous RPT significantly reduced the D50 metric, which measure the percentage of unique clonotypes accounting for 50% of total CDR3 sequences (Figure 4J). This shift toward dominant clonotypes demonstrates that absorbed dose heterogeneity from RPT drives infiltration of clonally expanded effector CD8 T cells within the TME.

### Improved clinical response in patients with high intratumoral dose heterogeneity from **^177^Lu-PSMA-617**

To assess clinical implications of these novel mechanistic findings on the immuno- radiobiological effects of RPT dose heterogeneity, we performed an exploratory analysis of all patients with metastatic prostate cancer who were treated at the University of Wisconsin with 177-lutetium vipivotide tetraxetan (^177^Lu-PSMA-617) and for whom multi-timepoint imaging-based dosimetry and ≥12 months of clinical follow-up was available (n=11, Figure 5A-B). We hypothesized that patients receiving highly heterogeneous intratumoral absorbed dose distribution, with at least one tumor receiving low-, moderate-, and high-dose in a single TME, would exhibit improved prostate-specific antigen (PSA) response and biochemical progression-free survival. Based on our preclinical mechanisms, we hypothesized that that high <u>intra</u>tumoral dose heterogeneity within a single tumor site could promote effector T cell clonal expansion, whereas <u>inter</u>tumoral heterogeneity would not suffice to achieve this same effect.

**Figure 5.**
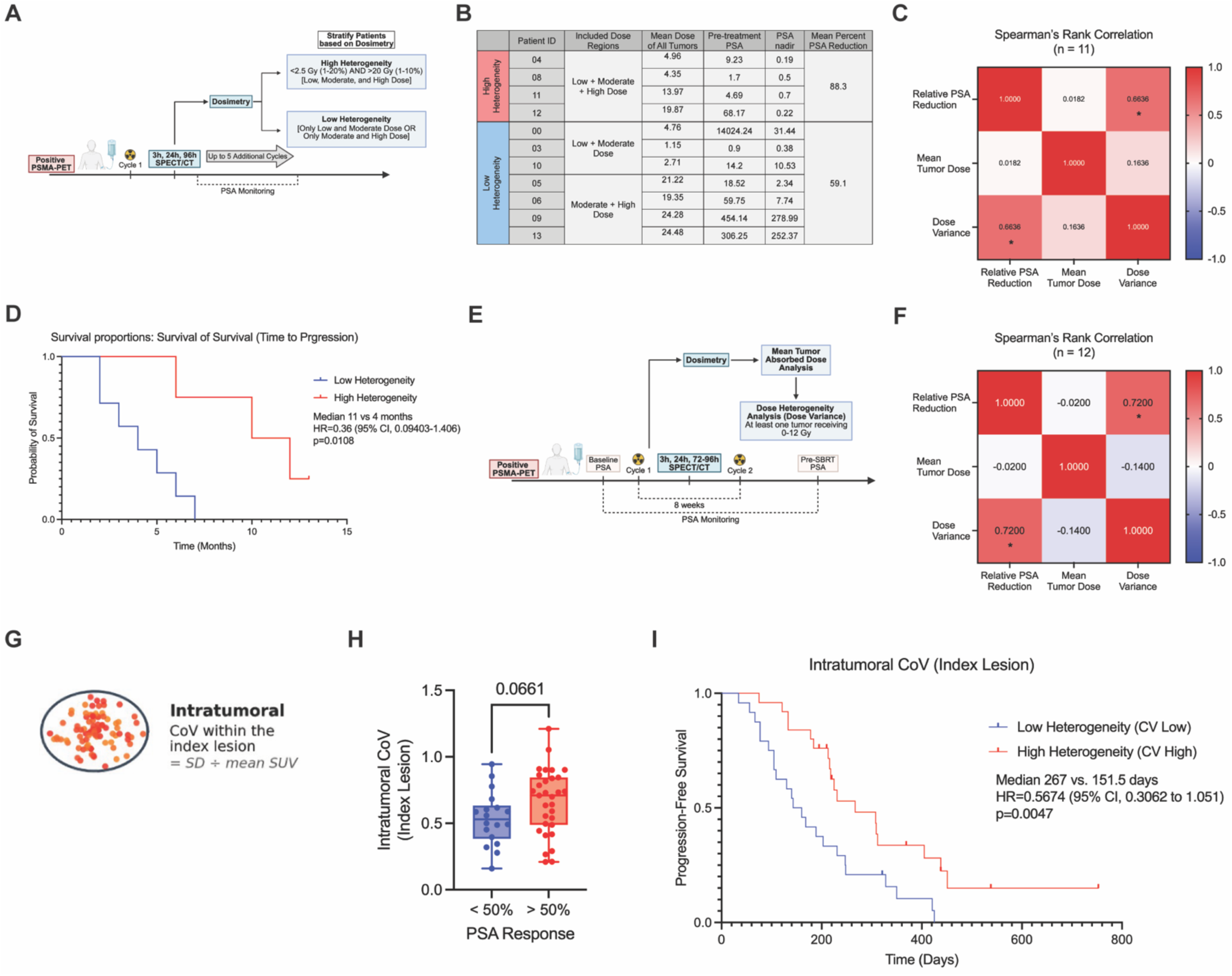
Intratumoral heterogeneity correlates to improved clinical response to ^177^Lu-PSMA-617 and ^177^Lu- PSMA-PNT2002. A) Schematic for ^177^Lu-PSMA-617 administration and imaging timing. B) Pre-treatment PSA and PSA nadir of stratified patients (heterogeneous dose distribution or homogenous dose distribution) and average percent reduction of PSA for the two groups (n = 12). C) Spearman’s rank correlation between relative PSA reduction, mean tumor dose, and dose variance (n = 12). D) Progression-free survival comparing patients stratified to high heterogeneity and low heterogeneity. E) Schematic for ^177^Lu-PNT2002 administration and imaging timing from the LUNAR trial. F) Spearman’s rank correlation between Mean Tumor Dose, Percent PSA Reduction, and Dose Variance for subset of patients with high heterogeneity (n = 12). G) Schematic of intratumoral coefficient of variation (CV), the SD divided by the mean of voxel SUV within the dominant (index) lesion. H) Index-lesion (intratumoral) CV in PSA50 responders (n = 31) versus non-responders (n = 18); PSA50 response 79% vs 48% for patients above versus below the median CV. I) Progression-free survival comparing high CV (high heterogeneity, n = 24) and low CV (low heterogeneity, n = 25). *p<0.05 by unpaired Spearman’s rank correlation.

Patients underwent SPECT/CT imaging at three timepoints (3, 24, and 96 hours) following cycle 1 of ^177^Lu-PSMA-617. Tumor segmentation was CT-based, and physician- contoured lesions underwent retrospective multi-timepoint dosimetry. Patient response was tracked for 12 months following the start of RPT treatment. Based on the preclinical dosimetry parameters (Figure 1C-1F), patients were stratified as high-heterogeneity if ≥1 lesion received ≤2.5 Gy to 1%-20% of the tumor volume and ≥20 Gy to 1%-10% of the tumor volume. Four of eleven patients met high-heterogeneity criteria in at least one lesion (Patient 04 with 3 lesions; Patients 08, 11, 12 with 1 lesion; Figure S6A). In the low-heterogeneity group, Patients 00, 03, and 10 lacked high-dose regions, while Patients 05, 06, 09, and 13 lacked low-dose preservation (Figure S6B).

Based on the multi-timepoint imaging-based dosimetry, we stratified patients into two groups, high-heterogeneity and low-heterogeneity. High-heterogeneity was defined based on our preclinical dosimetry data (Figure 1C-1F) as any one tumor receiving 2.5 Gy or less to 1% to 20% of the tumor and 20 Gy or more to 1% to 10% of the tumor. A patient was stratified to the high-heterogeneity group if they had at least one lesion that met these dose distribution parameters. Patients in the high-heterogeneity group demonstrated greater mean PSA reduction compared to the low-heterogeneity group (88.27% vs. 59.13%), despite receiving a lower mean tumor dose (10.8 Gy vs. 13.99 Gy; Figure 5B). Spearman rank correlation revealed a strong, significant correlation between intratumoral dose variance and relative (%) PSA reduction (r_s_ = 0.66, p = 0.031; Figure 5C), whereas mean tumor dose showed no correlation (r_s_ = 0.0200, p = 0.967). Patients in the high-heterogeneity group also experienced significantly prolonged biochemical progression-free survival (12 months vs. 4 months, p=0.0108; Figure 5D and S6C).

To test whether intertumoral dose heterogeneity could yield similar benefits, we evaluated Patients 09 and 13, who had separate tumors receiving distinct low, moderate, and high doses. The four patients exhibiting intratumoral dose heterogeneity demonstrated superior progression-free survival compared to the two patients with intertumoral heterogeneity (Figure S6D). While the sample size is small, these results are consistent with the hypothesis that a non-uniform distribution of RPT that includes low, moderate, and high dose regions in a single TME may improve response to RPT delivering lutetium-177.

### Validation of improved clinical response in a separate cohort of patients with high intratumoral dose heterogeneity from ^177^Lu-PSMA-PNT2002

To validate these findings in an independent cohort, we analyzed patients with oligo- recurrent hormone-sensitive prostate cancer from the randomized phase II LUNAR trial (NCT05496959), evaluating neo-adjuvant lutetium Lu 177 zadavotide guraxetan (^177^Lu-PSMA- PNT2002) followed by stereotactic body radiotherapy (SBRT) versus SBRT alone (40, 41). We evaluated the investigational arm subset (n=45) who underwent SPECT/CT imaging at three timepoints (3, 24, and 72-96 hours) following cycle 1 of ^177^Lu-PSMA-PNT2002 (Figure 5E). To avoid confounding effects from SBRT, PSA responses were analyzed from RPT initiation until immediately prior to SBRT delivery.

Mirroring our approach above, we evaluated the potential correlation between mean tumor dose or dose heterogeneity and relative PSA reduction. Due to lower tumor burden and smaller lesion size in the oligometastatic cohort, most patients did not receive moderate or high doses to any tumor region (n=33). In a subset analysis of patients receiving ≥12 Gy to any portion of at least one tumor site (n=12), Spearman rank correlation demonstrated a strong, significant correlation between intratumoral dose variance and relative PSA reduction (r_s_ = 0.72, p = 0.011; Figure 5F), whereas mean tumor dose did not correlate (r_s_ = -0.0200, p = 0.956). Across the full cohort (n=45), where most tumors lacked high-dose regions, neither mean absorbed dose nor dose variance correlated with PSA reduction (Figure S6E). These results support the hypothesis that heterogeneous distribution of RPT that delivers a range from low to high dose within at least one TME may play a critical role in promoting therapeutic response.

### Identification of patients with improved clinical response following ^177^Lu-PSMA-617 using AI-based assessment of heterogeneity in PSMA PET imaging

Because multi-timepoint SPECT-based dosimetry is resource intensive, we investigated whether baseline pre-treatment PSMA PET intratumoral heterogeneity could serve as a predictive radiographic biomarker in 49 patients with metastatic castrate-resistant prostate cancer treated with ^177^Lu-PSMA-617 at the University of Wisconsin Hospitals and Clinics. Baseline PSMA PET/CT index lesions (defined as the highest SUV_total_ tumor corresponding to a discrete CT lesion) were quantified using TRAQinformIQ (AIQ Solutions) (42). Intratumoral heterogeneity was defined by the coefficient of variation (CV) of the index lesion (Figure 5G).

When evaluating biochemical response, patients achieving ≥50% PSA reduction (responders) showed a strong trend toward higher baseline index lesion CV compared to non-responders (79% vs. 48%, p=0.066; Figure 5H). Furthermore, when the cohort was stratified by median index lesion CV into high-heterogeneity (n=24) and low-heterogeneity (n=25) groups, patients with higher baseline index lesion heterogeneity achieved significantly longer progression-free survival compared to those with lower heterogeneity (p=0.0047; Figure 5I). <u>Inter</u>tumoral SUV heterogeneity did not impact progression-free survival (Figure S5F). Our findings suggest that pre-treatment PET intratumoral heterogeneity at any one tumor site, but not intertumoral heterogeneity between lesions, may be an effective predictive radiographic biomarker for identifying patients who achieve greatest benefit from RPT.

## DISCUSSION

Our findings challenge a central assumption of conventional radiobiology: that optimal radiopharmaceutical therapy (RPT) requires uniform radiation coverage across all tumor sites. Instead, we demonstrate dose heterogeneity, delivering a spectrum from low (< 2.5 Gy) to high (> 20 Gy) absorbed doses within a single TME, while not inducing lymphopenia acts as a potent driver of adaptive anti-tumor immunity. Across two distinct syngeneic murine tumor models, heterogeneous lutetium-177 delivery induced dose-dependent expression of endothelial adhesion marker (*Icam*), a type I interferon response (*Ifnβ1*), and immune susceptibility marker (*H2-K1)* ultimately enhancing T cell-dependent survival compared to uniform dose distributions.

Importantly, this concept translates directly to human clinical response across three independent cohorts of men with metastatic prostate cancer. First in a retrospective dosimetry cohort, patients exhibiting high intratumoral dose heterogeneity (at least one lesion with low, moderate, and high dose regions) following ^177^Lu-PSMA-617 significantly correlated with PSA reductions and was associated with significantly prolonged biochemical progression-free survival (bPFS). Second, analysis of the prospective LUNAR trial revealed that while mean tumor absorbed dose did not correlate with relative PSA reduction, higher intratumoral dose variance strongly correlated with relative PSA reduction. Third, pre-treatment PSMA PET imaging demonstrated that upfront radiographic target heterogeneity independently predicts progression-free survival. Together, these preclinical and clinical data demonstrate that dose heterogeneity, rather than limiting efficacy, actively enhances RPT response by stimulating adaptive anti-tumor immunity.

Our clinical observations highlight a critical distinction between multi-timepoint dosimetry and single timepoint PET imaging. In contrast with our dosimetry-based analyses, assessment of single timepoint pre-treatment PSMA PET demonstrated a strong trend, but not a statistically significant correlation, with relative PSA reduction. This discrepancy likely results from the additional spatial and temporal information captured by multi-timepoint dosimetry relative to single timepoint PET imaging. Multi-timepoint dosimetry provides a more accurate estimate of delivered intratumoral dose heterogeneity, rendering it a more sensitive, though labor-intensive, approach for predicting this near-term biochemical response. Nevertheless, pre-treatment PET-based heterogeneity assessment was sufficient for identifying patient cohorts with improved progression-free survival, serving as an accessible predictive biomarker when serial SPECT/CT imaging and dosimetry are clinically impractical.

Our results identify potential immune-related mechanisms for the enhanced therapeutic response observed with heterogeneous RPT. TDLNs are critical for initiating and sustaining an anti-tumor immune response. Recent work demonstrated that TDLN-sparing RT led to enhanced anti-tumor immunity, whereas direct irradiation of the TDLN resulted in a loss of that response (43, 44). Furthermore, evidence from heterogeneous brachytherapy showed that delivering heterogeneous doses that include low-dose regions spares DC migration within the TDLN (9) and that low-dose RT has been shown to promote T cell clonal expansion, a process requiring direct T cell and DC interaction (31). Heterogeneous EBRT and brachytherapy have both also been shown to promote subsequent immune cell infiltration compared to uniform-field EBRT (9). The use of heterogeneous RT may deliver a high dose to some regions of the tumor stimulating the release of cytokines and DAMPs that drive immune cell infiltration into neighboring low-dose regions. Additionally, these low-dose regions may be critical to propagating an existing immune response or immune functionality within a tumor by sparing resident immune cells and/or their function. Indeed, we found that non-uniform RPT promoted DC activation in the TDLNs, preserved and recruited intratumoral CD8 T cells, and stimulated clonal expansion of T cell clonotypes within the TME.

These findings provide an actionable framework for personalizing RPT dosing and patient selection. Currently, RPT agents like ^177^Lu-PSMA-617 are prescribed at a fixed injected activity (45). With fixed activity dosing, mean absorbed doses to the tumors and bone marrow vary widely between individual patients (45). Individualized dosimetry can be used to overcome this challenge, allowing selection and dosing of patients in whom lymphopenia can be avoided by delivering a mean bone marrow dose under 2 Gy while delivering a maximal mean tumor dose that achieves a distribution of low- to high-dose radiation in the TME. Our results suggest that RPT dosing should be tailored to the therapeutic objective. Specifically, radiation-sensitive or poorly immunogenic tumors may benefit from maximizing mean dose for direct cytotoxicity, whereas radiation-resistant or highly immunogenic tumors, particularly when combined with immunotherapies like immune checkpoint inhibitors, may be most effectively treated by optimizing dose heterogeneity. Although heterogeneous target expression is conventionally regarded as a major barrier or mechanism of drug resistance, our findings demonstrate that RPT can leverage this spatial variation to generate immunogenic dose gradients. Furthermore, radionuclide selection plays a key role; shorter-range emissions (e.g., alpha particles) more readily preserve low-dose regions within the TME, permitting dose escalation and potentially achieving greater immunogenic effect.

We acknowledge several limitations to our study. Preclinically, while syngeneic murine tumor models are essential for evaluating intact adaptive immunity, they do not fully mirror human tumorigenesis, metastatic potential, or the tumor-immune microenvironment. Additionally, to control dose heterogeneity in the preclinical studies, we employed an exogenous FAP-targeting model (^177^Lu-PNT6555) to deliver heterogeneous radiation to the tumor cells rather than native stromal components; we are not able to comment whether the same results would be achieved if using RPT agents targeting stromal components of a tumor. Clinically, our retrospective discovery cohort was limited in sample size, and cross-trial analysis of the prospective LUNAR trial introduced technical constraints due to variations in SPECT/CT reconstruction protocols. In particular, limited spatial resolution in small, low-volume metastatic lesions can lead to partial-volume effects that potentially underestimate absolute intratumoral dose variance. Furthermore, while pre-treatment PSMA PET heterogeneity was sufficient for identifying cohorts with significantly improved progression-free survival, we acknowledge that a static, upfront scan cannot capture the dynamic tumor therapeutic interactions, such as target saturation or tracer clearance, that are directly measured by post-treatment dosimetry. Nevertheless, these independent clinical cohorts support a reconceptualization of how RPT dose and dose heterogeneity can be optimized to achieve therapeutic efficacy.

RPT dose heterogeneity has traditionally been viewed as a limitation that risks leaving portions of a tumor under-dosed. However, by leveraging radiation dose-dependent immune responses, heterogeneous RPT promotes a robust adaptive anti-tumor immune response that provides therapeutic benefit in the metastatic setting. Delivering a range of low to high dose in a single TME enhances systemic anti-tumor immunity while preserving immune cell function. This paradigm shift in immuno-radiobiology highlights the need for further research to explore and harness non-uniform distributions as an optimizable variable to enhance treatment efficacy and patient selection.

## METHODS

### Cell lines and Cell culture

The murine melanoma B16-F10 and B16-F10-mFAP cell lines were obtained from Tufts University. The murine KP sarcoma cell line, derived from the transgenic KP sarcoma models as previously described (46), was obtained from Dr. David Kirsch (University Health Network Princess Margaret Cancer Centre). B16-F10 and cell lines were grown in RPMI supplemented with 10% FBS, 100 U/mL penicillin, and 100 μg/mL streptomycin; KP sarcoma cells were grown in DMEM supplemented with 10% FBS, 100 U/mL penicillin, and 100 μg/mL streptomycin. Cells were incubated in a humidified incubator at 37°C with 5% CO_2_. Cell line authentication was performed per ATCC guidelines using morphology, growth curves, and Mycoplasma testing within 6 months of use.

### Murine tumor models

Mice were housed and treated under an Institutional Animal Care and Use Committee (IACUC) approved protocol (protocol number M005670) at the University of Wisconsin – Madison. Female C57BL/6 mice were purchased at age 6 to 8 weeks from Taconic. Female SvJ mice were purchased at age 6 to 8 weeks from Jackson Laboratory. B16-F10 tumors were engrafted by subcutaneous flank injection of 2 x 10^5^ tumor cells. KP Sarcoma tumors were engrafted by orthotopic gastrocnemius injection of 5 x 10^4^ tumor cells. Tumor size was determined using digital calipers and volume approximated as (length x width^2^)/2. Mice were randomized immediately before treatment by stratified randomization by tumor size. Only mice with palpable tumors the day before treatment began were included in the study. Treatment began when tumors were well-established, approximately 9 days after tumor implantation for B16-F10 (∼30-50 mm^3^) and KP sarcoma (∼30-50 mm^3^). The day of RPT was defined as “day 0” of treatment. Mice were euthanized when tumor size exceeded 20 mm in longest dimension or recommended by an independent animal health monitor for morbidity of moribund behavior. Mice bearing KP sarcoma tumors were monitored for cancer progression morbidity behavior to determine endpoint. Due to the institutional radiation safety protocols and the mice being treated with radioactive isotopes, all investigators were aware of mouse treatment groups. The number of mice in therapy groups was based on expected outcomes and an effort to power experimental design to anticipated effect size, determined in collaboration with biostatisticians, and therefore may vary between treatment groups and tumor models. Mouse therapy experiments were repeated in duplicate. Final replicates are presented for tumor response and aggregate data for survival; number of animals per group is indicated in figure legends.

### CD8+ depletion studies

CD8+ T cell depletion was performed by injection of anti-murine CD8α (Rat IgG2b, κ; Clone 2.43; BioXCell) or IgG2b control (Rat IgG2b, κ; Clone LTF-2; BioXCell) by 300 μg intraperitoneal injection on days 0, 5, and 10. Depletion was confirmed on treatment day 7 by flow cytometry.

### Radionuclides and Radiochemistry

^177^Lu was purchased as ^177^LuCl_3_ from SHINE Technologies. Radiochemistry was performed as previously described (25, 26).

### SPECT/CT imaging

Mice bearing subcutaneous B16-F10 (C57BL/6) or intramuscular KP sarcoma (SvJ) tumors were administered 500 μCi of ^177^Lu-PNT6555 or ^177^Lu-NM600 in the lateral tail vein, respectively. Individual mice (n=4) were placed prone into a MILabs U-SPECT6/CTUhr system (Houten, The Netherlands) under 2% isoflurane for longitudinal scans at 3, 24, 96, and 168 h post-injection. CT scans (5 min) were acquired for anatomical reference and attenuation correction and fused with the SPECT scans (45 min). Image reconstruction used a similarity-regulated ordered-subset expectation maximization (SROSEM) algorithm. Images were quantitatively analyzed by drawing volumes of interest (VOI) over the tumor and organs of interest to determine the percent injected activity (IA) per gram (%IA/g) for each tissue. An *ex vivo* biodistribution study was carried out after the last scan time point.

### iQID imaging

Mice bearing subcutaneous B16-mFAP^+^ and mixed B16-mFAP^+^/mFAP^-^ (C57BL/6) tumors were administered 666 μCi ^177^Lu-PNT6555 in the lateral tail vein. Mice bearing intramuscular KP sarcoma (SvJ) were administered 250 μCi ^177^Lu-NM600 in the lateral tail vein. Mice were euthanized at 24h post-injection and tumors were harvested, embedded in optimal cutting temperature (OCT) gel and frozen on dry ice. The tumor containing OCT blocks were cut at -16°C at a 20 μm thickness. The slide was placed on the iQID camera and imaged for 2h. iQID scans were acquired for the sections as count rate images, which were subsequently converted into activity concentration maps (Bq/mL) using a previously established calibration curve.

### *In vivo* and *ex vivo* dosimetry estimation

Macroscopic (∼mm) *in vivo* SPECT/CT-based ^177^Lu-PNT6555 dosimetry was performed according to a previously described method (27). Organ activities were quantified from ROIs delineated at each time point. Absorbed dose rates were derived from image-based Monte Carlo simulations and integrated using the trapezoidal method, including physical decay after the final time point, to determine the mean total absorbed dose.

For miscroscale (∼μm) *ex vivo* dosimetry, the absorbed dose rate of tumor sections at the time of harvesting was estimated using ionizing-radiation Quantum Imaging Detector (iQID) activity image-based Monte Carlo simulations in a tissue-equivalent medium (liquid water). This calculation considered only the self-dose within the individual tumor section, excluding cross-dose contributions from adjacent sections or organs. To calculate the mean total absorbed dose, this single time point dose rate was integrated by applying the specific pharmacokinetic parameters from the corresponding tumor model’s serial *in vivo* SPECT/CT data.

### Radiation therapy (RT)

^177^Lu-PNT6555 or ^177^Lu-NM600 therapy was administered via intravenous tail vein injection on treatment day 0 for B16 or KP sarcoma tumor-bearing mice. Table 1 shows injected activity for each RPT dose studied.

### Gene Expression analysis

Tumors were homogenized in TRIzol using a Bead Mill Homogenizer (Bead Ruptor Elite, Omni International). Total RNA was extracted using RNeasy Mini Kit (QIAGEN, Germany) followed by cDNA synthesis using QuantiTect Reverse Transcription Kit (QIAGEN, Germany) according to the manufacturers’ instructions. Quantitative polymerase chain reaction (qRT-PCR) was performed using TaqMan^TM^ Fast Advanced Master Mix. Thermal cycling conditions (QuantStudio 6, Applied Biosystems) included the UNG activation at 50°C for 2 min, followed by polymerase activation stage at 95°C for 2 min followed by 40 cycles of each PCR step (denaturation) 95°C for 1s and (annealing/extension) 60°C for 20s. Fold change normalized to untreated control samples was calculated using the ΔΔCt method in Excel with Rn18S as an endogenous control. A complete list of TaqMan probes is provided in Table S1.

### Lymphopenia Assessment

To evaluate ^177^Lu-NM600 effect on lymphopenia, complete blood count (CBC) studies were performed in therapy mice. Groups of KP sarcoma tumor bearing mice (n=10/group) were treated with 0, 2.5, 5, 10, and 15 Gy from ^177^Lu-NM600 intravenously. Submandibular blood collection was performed on five mice per treatment group at day 7 and 14 post-injection. CBC analysis was completed using whole blood and an Abaxis VetScan HM5 hematology analyzer (Union City, CA).

### Flow cytometry

Flow cytometry was performed as previously described (47), using fluorescent beads (UltraComp Beads eBeads, 176 Invitrogen) to determine compensation, and fluorescence minus one (FMO) methodology to determine gating. Rainbow beads (Spherotech) were used to set voltages for each flow cytometry timepoint. For *in vivo* analysis, tumors and TDLN were harvested and manually dissociated. Blood was collected by submandibular vein collection, treated with RBC lysis buffer (BioLegend), and washed with phosphate-buffered saline prior to staining. Cells were treated with CD16/32 antibody (BioLegend) to prevent non-specific binding. Live cell staining was performed using Ghost Red Dye 780 (Tonbo Biosciences) according to manufacturer’s instruction. After live-dead staining, a single cell suspension was labeled with the surface antibodies at 4°C for 30 min and washed three times using filtered flow buffer (2% FBS + 2 mM EDTA in PBS). For intracellular staining, cells were fixed and stained for internal markers with permeabilization solution according to manufacturer’s instructions (BD Cytek FOXP3 / Transcription Factor Staining Buffer Kit). Flow cytometry was performed using an Attune NxT Flow Cytometer (ThermoFisher). Data was analyzed using FlowJo Software. A complete list of antibody targets, clones, and fluorophores is provided in **Table S2**. Number of animals per group is indicated in figure legends.

### TCR sequencing

On day 12 after RPT injection, B16-mFAP^+^ and mixed B16-mFAP^+^/mFAP^-^ tumors were harvested from mice treated with non-radiolabeled PNT6555 or 5 Gy from ^177^Lu-PNT6555. Tumors were homogenized in TRIzol using a mortar and pestle. Total RNA was isolated as previously described followed by sample library preparation with the SMARTer Mouse TCR α/β Profiling Kit (634402, Takara). Pooled libraries were sequenced with miSeq (Illumina). The Alpha and Beta clonotype chains were assembled using MiXCR (48).

### Dosimetry of ^177^Lu-PSMA-617

Dosimetry was performed under UW24007. Dosimetry for patients undergoing ^177^Lu- PSMA-617 therapy was performed using multi-timepoint SPECT/CT performed Day 0 (3 ± 2 h), Day 1 (24 ± 4 h), and Day 4 (96 ± 24h) post ^177^Lu-PSMA-617 infusion. The datasets were deformably registered using the MIM Vista software. Tumors and normal organs were segmented and approved by the attending physician. The registered images and structure sets were imported into the Torch dose calculation software. Here, pharmacokinetic modeling was performed for each tumor and normal organ to generate time activity curves, which were integrated to obtain the cumulated activity. A voxel level Monte Carlo simulation was performed to obtain the dose distribution throughout the patient.

### Dosimetry of ^177^Lu-PSMA-PNT2002

Dosimetry was performed as part of the LUNAR trial (NCT05496959). Dosimetry for patients undergoing ^177^Lu-PSMA-PNT2002 therapy was performed using multi-timepoint SPECT/CT performed at Day 0 (4.3 ± 0.5 h), Day 1 (23.9 ± 1.2 h), and Day 3\4 (81.0 ± 11.8 h) post ^177^Lu-PSMA-PNT2002 infusion. Reconstruction of SPECT images was performed using the MIM SPECTRA Quant with 48 iterations, 1 subset, and no post filter applied. Individual tumor segmentation alignment between timepoints were manually verified to generate time activity data which was then fit with a monoexponential function and voxel S value applied. A phantom derived volume-dependent partial volume correction applied to determine an absorbed dose (41).

### Analysis of PSMA PET

Baseline diagnostic PSMA PET/CT was acquired with either [⁶⁸Ga]Ga-PSMA-11 or [¹⁸F]DCFPyL/piflufolastat. Lesion-level quantification was performed with TRAQinformIQ (AIQ Solutions), that automatically detects, segments, and characterizes disease ROIs and matches PET findings to CT correlates (42). The index lesion was defined as the ROI with the highest SUV_total_ corresponding to a discrete CT-identifiable lesion. Intratumoral heterogeneity was quantified using the coefficient of variation (CV = standard deviation_SUV/_mean_SUV_) of the index lesion, and patients were stratified into high- and low-heterogeneity cohorts using median split analysis.

## Statistical analysis

Prism 10 (GraphPad Software) was used to create figures and calculate t-tests, ANOVA, and Spearman’s rank correlation. Student’s t-test was used for two-group comparisons. Two-way ANOVA with Tukey’s honestly significant difference (HSD) test to adjust for multiple comparisons was used to assess statistical significance of observed mean differences in gene expression and flow cytometry. Spearman’s rank correlation was used to assess the correlation over two dependent, overlapping groups. R software (version 4.4.1) was used for tumor growth analysis; all available data was used. To compare treatment groups, linear mixed models after log base 10 transformation of tumor volume were fitted on cell line, time in days, and their interaction using all available data. Pairwise contrasts were adjusted using Tukey’s method. Kaplan-Meier method was used to estimate the survival distribution for the overall survival. Then, pairwise comparison of the overall survival was made using a log-rank Benjamini-Hochberg adjustment of p-values between levels of factors. All data are reported as mean ± standard error of the mean (SEM) unless otherwise noted. For all graphs, *, P < 0.05; **, P < 0.01; ***, P < 0.001; and ****, P < 0.0001.

## Acknowledgements

The authors’ work is supported in part by grants from NIH NCI P01CA250972, NIH NCI P50DE026787, UWCCC Support Grant NIH NCI P30CA014520, University of Wisconsin Small Animal Imaging & Radiotherapy Facility, and NIH S10OD028670-01, UWCCC Flow Cytometry Laboratory. The authors would like to thank the University of Wisconsin Carbone Cancer Center (UWCCC) for supporting this project. The authors would like to thank Dr. Willliam Bachovchin from Tufts university for providing the FAP engineered cell lines. The authors would like to thank Point Biopharma Eli Lily and Archeus for providing key radiopharmaceutical agents.

## Author Contribution

MET, CS, MA, TS, PAC: animal work and analysis. MET, WJJ, PAC, TC: flow cytometry staining and analysis. SHA, APB, MBI, RTH: radiochemistry. MET, TC, RK: RPT biodistribution and toxicity studies. MET, WWJ, CS: qPCR analysis and sample preparation. MET and YW: TCR sequencing sample preparation. OK, CDH, BPB: SPECT analysis and ^177^Lu-PNT6555 and ^177^Lu-NM600 dosimetry. AOA, OK, BPB: iQID acquisition and analysis. MET and MH: biostatistical analysis. RWS and IO: bioinformatics analysis. MET, ML, AB, JF, TB, SP: Patient SPECT analysis, dosimetry analysis, and clinical response data for ^177^Lu-PSMA-617. ZE, CM, AUK, JC: Patient SPECT analysis, dosimetry analysis and clinical response data for ^177^Lu-PSMA-PNT2002. MET and ZSM: experimental design. JW, ZSM: funding, planning, and supervision of the project and data interpretation. MET, TB, ZSM wrote the manuscript. All authors read and approved the final manuscript.

