## Supplemental Data for "Intratumoral dose heterogeneity promotes adaptive anti-tumor immunity and predicts clinical response to radiopharmaceutical therapy"

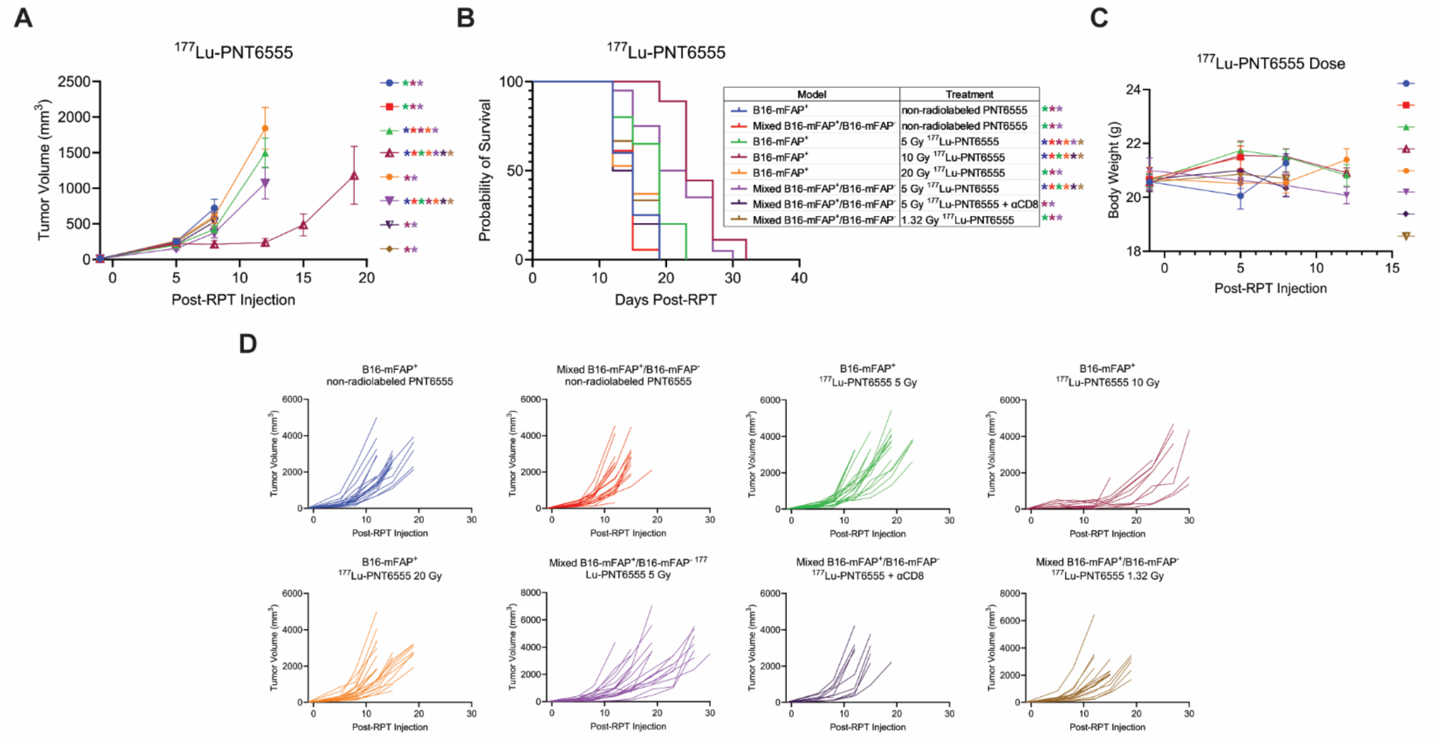

**Supplementary Figure S1. Overall tumor growth, survival, and body weight for $^{177}$Lu-PNT6555.** B16-mFAP$^{+}$ or mixed B16-mFAP$^{+}$/mFAP$^{-}$ tumor-bearing mice were treated with non-radiolabeled PNT6555,1.32, 5, 10, 20 Gy $^{177}$Lu-PNT6555. A) Overall tumor growth. B) Overall survival. C) Overall mouse body weights. D) Individual tumor growths for each treatment group. *p < 0.05 by linear mixed model (A), by Kaplan-Meier method (B) the color of the asterisk represents the group from which the group differs.

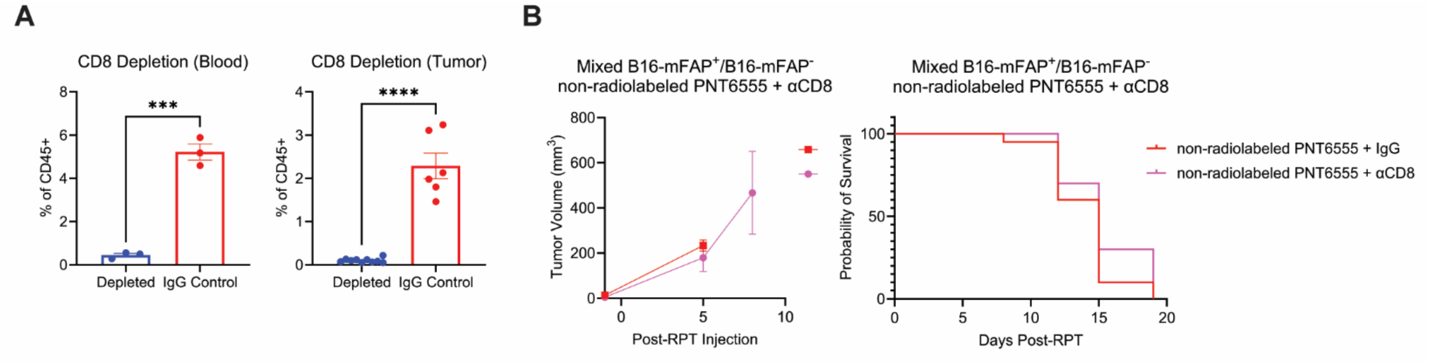


**Supplementary Figure S2. CD8 depletion alone does not affect tumor growth or survival.** B16-mFAP$^+$ or mixed B16-mFAP$^+$/mFAP$^-$ tumor-bearing mice were treated with control IgG or anti-CD8 (αCD8) antibodies. A) Confirmation of CD8 T cell depletion by flow cytometry on peripheral blood and flank tumor of mice. N=3-10 per group. ***$p < 0.001$. ****$p < 0.0001$ by T-test. B) Tumor growth curves and survival plots are shown. N=8 mice per group.

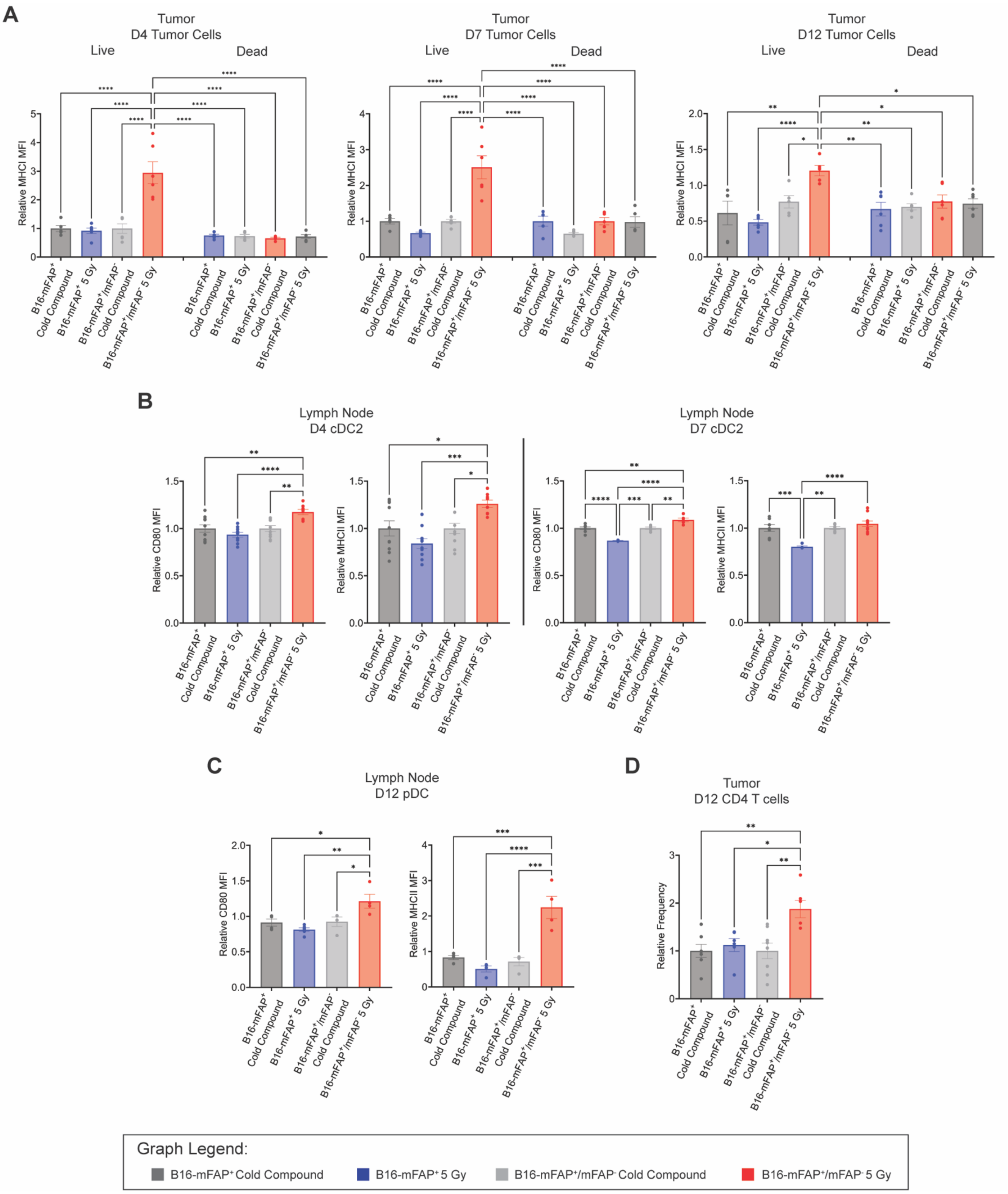
A
Tumor
D4 Tumor Cells
Live
Dead
Relative MHCI MFI
Tumor
D7 Tumor Cells
Tumor
D12 Tumor Cells
B16-mFAP+ Cold Compound
B16-mFAP+ 5 Gy
B16-mFAP+/mFAP- Cold Compound
B16-mFAP+/mFAP- 5 Gy
B
Lymph Node
D4 cDC2
Relative CD80 MFI
Relative MHCII MFI
Lymph Node
D7 cDC2
C
Lymph Node
D12 pDC
D
Tumor
D12 CD4 T cells
Relative Frequency
Graph Legend:
B16-mFAP+ Cold Compound
B16-mFAP+ 5 Gy
B16-mFAP+/mFAP- Cold Compound
B16-mFAP+/mFAP- 5 Gy

**Supplementary Figure S3. Radiopharmaceutical therapy dose heterogeneity including a low, moderate, and high dose region leads to increased MHCI expression and T cell infiltration in the tumor and dendritic cell activation in the tumor draining lymph node.** A-D) B16-mFAP$^+$ or mixed B16-mFAP$^+$/mFAP$^-$ tumor-bearing mice were treated with non-radiolabeled PNT6555 (Cold Compound) or 5 Gy $^{177}$Lu-PNT6555 on day 0 and tumors and tumor draining lymph nodes were harvested at day 4, 7, and 12. Results shown as relative frequencies and MFIs to non-radiolabeled PNT6555 (Cold Compound). A) Analysis of MHCI expression on live and dead tumor cells (CD45-MHCI+) at day 4, 7, 12. B) Analysis of CD80 and MHC class II MFI in TDLN type 2 conventional dendritic cells, cDC2 (CD45+CD11b+CD11c+MHCII+CD172a+) at day 4 and 7. C) Analysis of CD80 and MHC class II MFI in TDLN plasmacytoid dendritic cells, pDC (CD45+CD11b+CD11c+B220+SiglecH+) at day 12. D) Analysis of CD4+ T cell (CD45+CD3+CD4+) infiltration in tumor at day 12. *$p < 0.05$. **$p < 0.01$, ***$p < 0.001$, ****$p < 0.0001$ by one-way ANOVA (A-D).

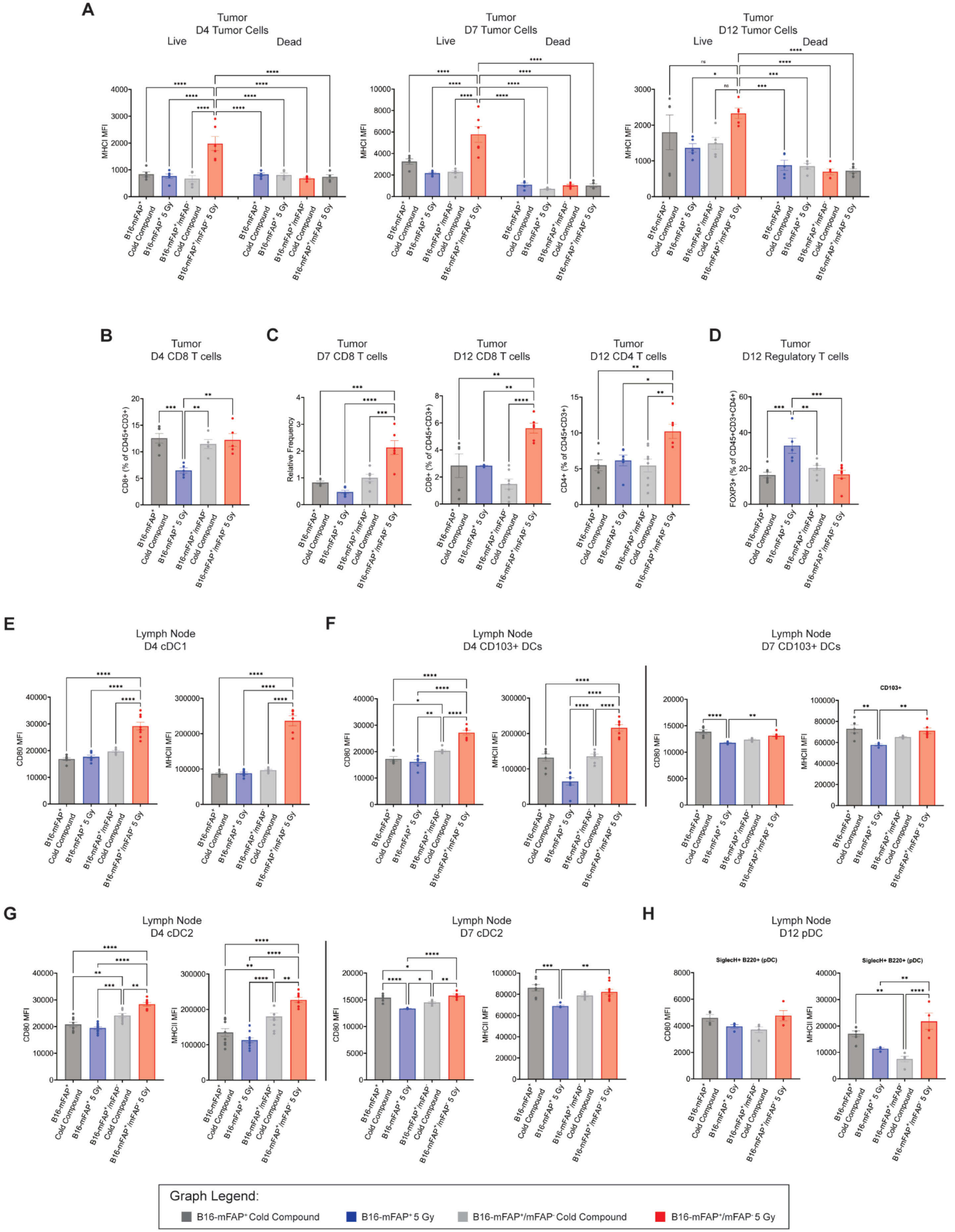
A
Tumor
D4 Tumor Cells
D7 Tumor Cells
D12 Tumor Cells
Live
Dead
MHCI MFI
B
Tumor
D4 CD8 T cells
CD8+ (% of CD45+CD3+)
C
D7 CD8 T cells
Relative Frequency
D12 CD8 T cells
D12 CD4 T cells
CD4+ (% of CD45+CD3+)
D
D12 Regulatory T cells
FOXP3+ (% of CD45+CD3+CD4+)
E
Lymph Node
D4 cDC1
CD80 MFI
MHCII MFI
F
D4 CD103+ DCs
D7 CD103+ DCs
CD103+
G
D4 cDC2
D7 cDC2
H
D12 pDC
SiglecH+ B220+ (pDC)
Graph Legend:
B16-mFAP+ Cold Compound
B16-mFAP+ 5 Gy
B16-mFAP+/mFAP- Cold Compound
B16-mFAP+/mFAP- 5 Gy

**Supplementary Figure S4. Raw MFIs for MHCI expression and dendritic cell activation and raw cell frequencies for CD8 and CD4 T cells.** A-I) B16-mFAP$^+$ (B16-mFAP) or mixed B16-mFAP$^+$/mFAP$^-$ tumor-bearing mice were treated with non-radiolabeled PNT6555 or 5 Gy $^{177}$Lu-PNT6555 on day 0 and tumors and tumor draining lymph nodes were harvested at day 4, 7, and 12; the raw values are shown for MHCI expression on live and dead tumor cells at day 4, 7, 12 (A), CD8$^+$ T cells preservation in tumor at day 4 (B), CD8+ T cell and CD4+ T cell infiltration in tumor at day 12 (C), Regulatory T cell infiltration in tumor at day 12 (D), CD80 and MHC class II MFI in tumor draining lymph node (TDLN) type 1 conventional dendritic cells, cDC1 at day 4 (E), CD80 and MHC class II MFI in TDLN CD103+ DCs at day 4 and 7 (F), CD80 and MHC class II MFI in TDLN type 2 conventional dendritic cells, cDC2 at day 4 and 7 (G), CD80 and MHC class II MFI in TDLN plasmacytoid dendritic cells, pDC at day 12 (H). One-way ANOVA was used to compare expression between groups.

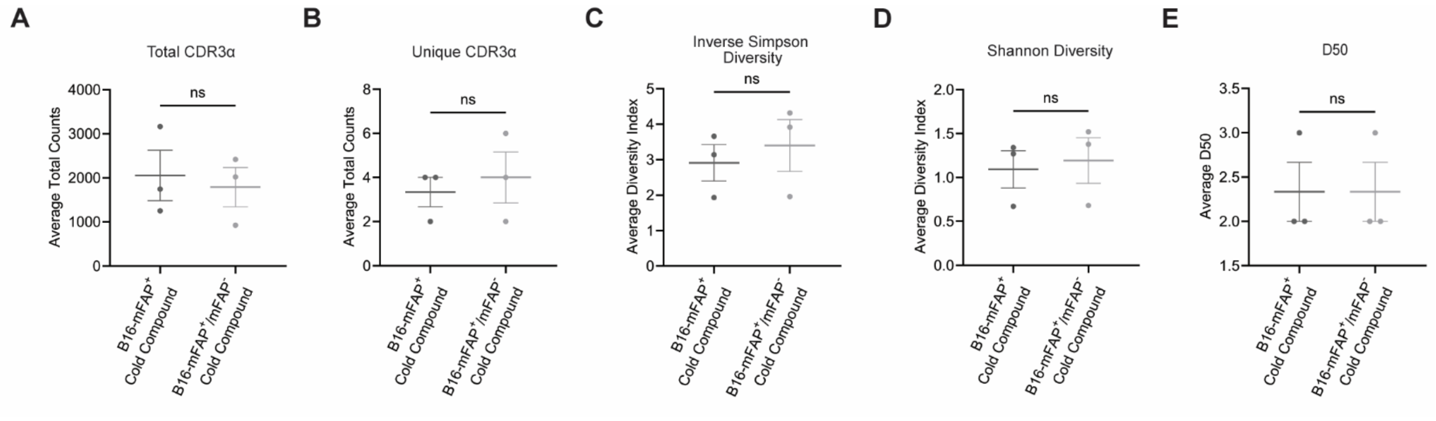


**Supplementary Figure S5. Target expression composition does not affect clonal expansion and TCR diversity.** A-E) TCR sequencing of RNA extracted from non-radiolabeled PNT6555 treated B16-mFAP$^{+}$ or mixed B16-mFAP$^{+}$/mFAP$^{-}$ tumor. A) Total number of CDR3α sequences. B) the counts of the unique CDR3α sequences. C) the inverse Simpson diversity index. D) the Shannon diversity index. E) the D50 value. n = 3 mice per group. Unpaired T-Test was used to compare the average CDR3α counts [(A) and (B)], diversity index [(C) and (D)], and D50 score (E) between groups.

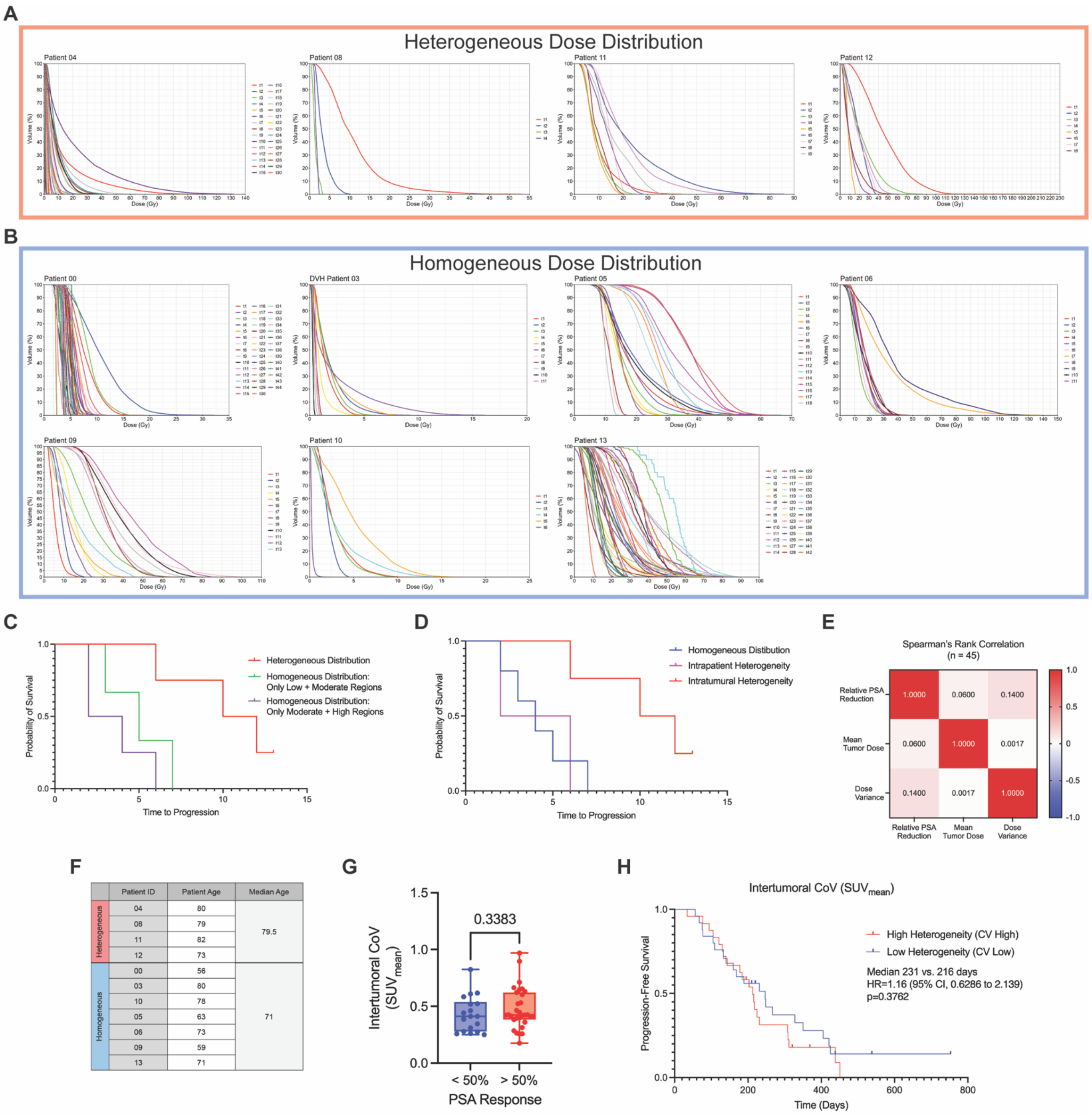


**Supplementary Figure S6. Cumulative DVH curves of all tumor lesions from patients treated with $^{177}$Lu-PSMA-617 and intertumoral progression-free survival.** A-B) Cumulative DVH curves for all contoured lesions from patients treated with $^{177}$Lu-PSMA-617 separated into the heterogeneous distribution group (red) and homogeneous distribution group (blue). C) Progression-free survival with homogeneous distribution separated into only low and moderate dose regions and only moderate and high dose regions. D) Progression-free survival comparing intratumoral dose heterogeneity to intertumoral dose heterogeneity. E) Patient age at start of treatment. F) Intertumoral CV in PSA50 responders versus non-responders. J) Progression-free survival comparing high intertumoral CV and low intertumoral CV.

**Table S1.** List of TaqMan probes

| Gene Name | Assay ID |
|---|---|
| *Icam1* | Mm00516023_m1 |
| *Ifnβ1* | Mm00439552_s1 |
| *H2-K1* | Mm04208017_mH |
| *Rn18s* | Mm04277571_s1 |

**Table S2.** List of flow cytometry antibody targets, clones and fluorophores

| Name | Clone | Fluorophore | Catalog number |
|---|---|---|---|
| CD3 | 17A2 | FITC | BioLegend 100204 |
| CD8a | S18018E | PerCP-eFluor710 | BioLegend 162310 |
| FOXP3 | MF-14 | PE | BioLegend 126404 |
| NK1.1 | S17016D | PE-Dazzle594 | BioLegend 156518 |
| CD45 | S18009F | PE-Cy7 | BioLegend 157206 |
| MHC-I (H-2K$^b$) | AF6-88.5 | BV421 | BioLegend 116525 |
| CD4 | GK1.5 | BV510 | BioLegend 100449 |
| CD11c | N418 | BV605 | BioLegend 117334 |
| B220 | RA3-6B2 | BV711 | BioLegend 103255 |
| CD25 | 3C7 | APC | BioLegend 101910 |
| CD11b | M1/70 | FITC | BioLegend 101206 |
| CD11c | N418 | PerCP-Cy5.5 | BioLegend 117328 |
| CD172a | P84 | PE | BioLegend 144012 |
| XCR1 | ZET | PE-Dazzle594 | BioLegend 148234 |
| CD45 | S18009F | PE-Cy7 | BioLegend 157206 |
| CD86 | PO3 | BV421 | BioLegend 105123 |
| MHCII (I-A/I-E) | M5/114.15.2 | BV510 | BioLegend 107636 |
| CD103 | 2E7 | BV605 | BioLegend 121433 |
| CD80 | 16-10A1 | BV711 | BioLegend 104743 |
| Siglec H | 551 | Alexa647 | BioLegend 129608 |
| B220 | RA3-6B2 | Alexa700 | BioLegend 103232 |
